# New highly selective antivirals for Chikungunya Virus identified from the screening of a drug-like compound library

**DOI:** 10.1101/2024.05.17.594707

**Authors:** Eliana F. Castro, Diego E. Álvarez

## Abstract

Chikungunya fever is a mosquito-borne disease caused by Chikungunya virus (CHIKV). Treatment of CHIKV infections is currently supportive and does not limit viral replication or symptoms of persistent chronic arthritis. Although there are multiple compounds reported as antivirals active against CHIKV *in vitro*, there are still no effective and safe antivirals. Thus, active research aims at the identification of new chemical structures with antiviral activity. Here, we report the screen of the Pandemic Response Box library of small molecules against a fully infectious CHIKV reporter virus. Our screening approach successfully identified previously reported CHIKV antiviral compounds within this library and further expanded potentially active hits, supporting the use of reporter-virus-based assays in high-throughput screening format as a reliable tool for antiviral drug discovery. Four molecules were identified as potential drug candidates against CHIKV: MMV1634402 (Brilacidin) and MMV102270 (Diphyllin), which were previously shown to present broad-spectrum antiviral activities, in addition to MMV1578574 (Eravacycline), and the antifungal MMV689401 (Fluopicolide), for which their antiviral potential is uncovered here.

## Introduction

Chikungunya fever is a mosquito-borne disease caused by Chikungunya virus (CHIKV), an alphavirus of the *Togaviridae* family. Alphaviruses are classified as arthritogenic alphaviruses, such as CHIKV, Ross River, Sindbis (SINV) and O’nyong-nyong, and encephalitic alphaviruses, such as Venezuelan (VEEV), Eastern (EEEV) and Western (WEEV) equine encephalitic viruses. Following the first CHIKV outbreak in 1952 in Tanzania, outbreaks were sporadic in Africa and Southeast Asia until 2004 when it reached the Western Hemisphere and CHIKV established as a global pathogen that continues to expand with the potential to generate significant outbreaks [1]. Initially, the virus diverged into three genotypes: West African (WA), East, Central and Southern Africa (ECSA), and Asian. In 2004, an epidemic of ECSA genotype started in Kenya and spread in 2005-2006 to the islands of the Indian Ocean, notably the Reunion Island (Pialoux 2007), giving rise to indian-ocean lineage (IOL). Since then, other significant outbreaks have occurred: the first Chikungunya outbreak in Europe was recorded in Italy in 2007, in the following years in France [2–4], and in 2014 the Asian genotype was introduced to the Caribbean, causing the first epidemic on the American continent [5]. Since 2017, an average of 166,000 cases have been reported annually in the Americas. Notably, in 2023 these cases amounted to 239,962 and 151,700 have already accumulated so far in 2024 [6, 7].

CHIKV transmission occurs from the bite of an infected *Aedes* mosquito, although in recent epidemics some cases resulted from maternal-fetal transmission [8]. CHIKV infection primarily causes musculoskeletal inflammation with debilitating symptoms, such as arthralgia, arthritis, and myalgia. Subclinical infections are rare (around 15%) and, in approximately 50% of infected patients, severe arthralgia and myalgia can last months or years [9]. Strategies to reduce CHIKV transmission and infection occur at the individual and community levels. Control of the mosquito vector is a strategy to control infections by CHIKV and other arboviruses. On the other hand, in November 2023, the FDA approved the attenuated virus vaccine Ixchiq for use in an emergency context for the prevention of CHIKV infections in adults in the United States [10, 11]. However, confirmatory trials to verify its clinical benefit are still required for its final approval. In turn, post-marketing studies should be conducted to evaluate the risk of serious chikungunya-like adverse reactions following the administration of Ixchiq. For the treatment of CHIKV infections, current supportive therapy to alleviate fever and pain uses paracetamol/acetaminophen, which does not limit viral replication or the persistence of chronic arthritis symptoms.

CHIKV has a positive polarity RNA genome of approximately 12 Kb encoding 4 non-structural proteins (nsp1-4) and 5 structural proteins (C, E3, E2, 6K, and E1). The surface of the virion contains 80 trimeric spikes formed by the glycoproteins E1 and E2. E2 facilitates the binding of the virus to its cellular receptor, after which the virus enters the host cell through a process of clathrin-mediated endocytosis. The fusion of the viral envelope with the endosome membrane is triggered by the decrease in pH in the vesicle that leads to conformational changes in the E1 glycoprotein that functions as a fusogen. The viral nucleocapsid is then released into the cytoplasm where uncoating of the viral genome occurs. Replication of the viral genome is preceded by its translation resulting in the production of the ns polyproteins P123 and P1234, which are processed by the cysteine-protease domain of nsp2 to produce the mature ns proteins. The nsp1 protein has NTPase, RNA triphosphatase, RNA helicase, and protease activity. Nsp3 interacts through its N-terminal macrodomain with negatively charged polymers including RNA molecules and its C-terminal domain, which is intrinsically disordered, would act as a binding platform for multiple cellular proteins that would be important for viral infection. Nsp4 of CHIKV is the viral RNA-dependent RNA polymerase. The glycoproteins that are part of the structural polyprotein are translated on membrane-associated ribosomes and are processed and post-translationally modified by cellular enzymes. Assembly and budding of virions occur at the plasma membrane of the infected cell (reviewed in [12]).

Although there are multiple compounds targeting different viral proteins reported as antivirals active against CHIKV *in vitro* [13, 14] there are still no effective and safe antivirals for CHIKV, making it necessary to search for new chemical structures.

Various international non-profit organizations offer large libraries of molecules for hit identification. For example, Medicines for Malaria Venture (MMV) [15] and the Drugs for Neglected Diseases initiative (DNDi) [16], are engaged in the research and development of new treatments for neglected diseases, both by repositioning or modifying drugs used in other therapeutic areas, and by identifying and developing entirely new chemical entities. The Pandemic Response Box (PRB), launched by the MMV and DNDi, provides 400 structurally diverse compounds that are already commercialized or in various stages of development [17, 18].

There is a previous report [19] where three compounds within the PRB were identified as antiviral hits against CHIKV following a subgenomic reporter-based primary screening, after which the antiviral activity of one of them (MMV690621 -GSK-983-) was confirmed. Here, we report the screen of the PRB library against a fully infectious CHIKV reporter virus. The approach allowed us to expand the number of hits in the primary and secondary screenings leading to the identification of four new molecules as potential antivirals against CHIKV with activity in the low micromolar range and excellent selectivity.

## Materials and methods

### Cells and Virus

Vero cells (ATCC CCL-81) were cultured at 37°C and 5% CO2 in complete Dulbecco’s modified Eagle’s medium (DMEM, Gibco) containing 10% fetal bovine serum (FBS) (Internegocios, Mercedes, Buenos Aires, Argentina), 100 IU/mL penicillin, and 100 mg/mL streptomycin (Gibco) (DMEM-10). For infections, Vero cells were cultured in DMEM supplemented with 2% FBS (DMEM-2).

CHIKV-ZsGreen were derived from infectious cDNA clones and reporter virus was obtained as previously reported [20, 21]. Viral titers of the final stocks were determined by fluorescent Foci Forming Units (FFU) assay. Viral stocks were stored at -70 °C until use.

### Compounds preparation

Pandemic Response Box (PRB) was kindly provided by Medicines for Malaria Venture (MMV) and the Drugs for Neglected Diseases Initiative (DNDi). For screening assays, compounds were provided in dry film form and 10μL of DMSO was added to reach a concentration of 10 mM (compounds 1-391) or 2 mM (compounds 392-400). For hits confirmation, solid samples of the selected compounds were provided and were dissolved in DMSO at 100 mM for cytotoxicity assays and from 2.5 to 0.004 mM (according to the compound) for antiviral assays. The compounds were then diluted in DMEM-2 to reach the desired concentration, maintaining 1% DMSO in each tested dilution.

### Primary screening

For primary screening, Vero cells were seeded in 96-well plates at a density of 1.5 x 10^4^ cells per well in DMEM-10 and incubated at 37°C and 5% CO_2_. After 24 h, the culture medium was removed, monolayers were washed twice with phosphate-buffered saline (PBS), infected with at least 700 FFUs of CHIKV-ZsGreen and simultaneously treated with 50 and 10 μM of the tested compounds in triplicate. Untreated infected cells (VC) and uninfected cells (CC) were included as controls. After 20 h at 37°C and 5% CO_2_, cultures were fixed with paraformaldehyde 4% at 4°C for 1 h. The fluorescent protein ZsGreen is expressed because of viral replication in infected cells and fluorescent infectious foci can be quantified. Automated counting of fluorescent infectious foci was performed using the EliSpot equipment and the InmunoSpot 7.0.30.2 software (ImmunoSpot C.T.L, Germany). Using this methodology, antiviral activity is detected as a decrease in the number of infectious foci respecting VC and dependent on the concentration of the compound. After foci quantification, fixed cultures were stained with crystal violet solution in methanol and plates were scanned for determination of media absorbance at 585 nm to estimate compound’s cytotoxicity respecting CC.

### Secondary Screening

In secondary screening, 36 compounds selected after primary screening were tested in five serial dilutions in triplicate (50-3.12 μM or 10-0.625 μM, depending on the compound tested). Antiviral assays were performed as described above. The focus count data for each antiviral concentration and the VC were used to plot dose-response curves that were fitted by nonlinear regression using GraphPad Prism 8 software. Effective concentration 50 (EC_50_) values, which are defined as the concentration of compound that reduces fFFU number by 50% with respect to VC, were calculated from dose-response curves. Cytotoxicity was evaluated by MTS/PMS method as described below.

### Cytotoxicity assays

To determine cytotoxicity, Vero cells were seeded in 96-well plates at a density of 1.5 x 10^4^ cells per well in DMEM-10 and incubated at 37°C and 5% CO_2_. After 24 h, the culture medium was removed, monolayers were washed twice with PBS, and serial dilutions of the test compounds were added. Cultures were incubated 24 h at 37°C, and then, cell viability was determined by the MTS/PMS method according to the manufacturer’s instructions (Promega). After 3 h at 37°C and 5% CO_2_, the absorbance was determined at 490 nm. Media absorbance obtained from each antiviral concentration and the CC were used to plot dose-response curves that were fitted by nonlinear regression using GraphPad Prism 8 software. The cytotoxic concentration 50 (CC_50_) values, which are defined as the concentration of compound that reduces absorbance_490_ by 50% with respect to CC, were estimated from dose-response curves. Finally, the EC_50_ and CC_50_ values were used to calculate the Selectivity Index (SI = CC_50_/EC_50_).

### Hit confirmation assays

Nine compounds were selected after secondary screening. To confirm their anti-CHIKV activity compounds were tested in six serial dilutions in triplicate against CHIKV-ZsGreen. Antiviral assays were performed as described above. Before fixing with paraformaldehyde 4%, culture supernatants from the VC and from the first, third and fifth tested concentration of each compound were collected and stored at -70ºC. Cytotoxicity was evaluated by MTS/PMS method as previously described. EC_50_ and CC_50_ values were obtained from three independent experiments and SI were calculated.

### Viral yield assays

To evaluate the effect of tested compounds on viral yield, culture supernatants collected from hits confirmation assays were titrated in Vero cells by FFU method. Briefly, Vero cells were seeded in 96-well plates at a density of 1.5 x 10^4^ cells per well in DMEM-10 and incubated at 37°C and 5% CO_2_. After 24 h, the culture medium was removed, monolayers were washed twice with PBS, and infected with 10; 20 or 40 μl of culture supernatant in a final volume of 50 μl. After 1 h at 37°C and 5% CO_2_, the culture medium was removed, monolayers were washed twice with PBS, and 100 μl of DMEM-2 was added per well. Cultures were incubated 20 h at 37°C and 5% CO_2_ and fixed with paraformaldehyde 4% at 4°C for 1 h. Automated numbering of fluorescent infectious foci was performed as described above and viral yield was estimated as fFFU/ml.

## Results

### Primary and secondary screening with CHIKV-ZsGreen identified nine active compounds within the PRB

To identify potentially active compounds against CHIKV, we carried out a reporter-based HTS antiviral assay using infectious CHIKV expressing ZsGreen (CHIKV-ZsGreen) under the control of the virus subgenomic promoter. In cells infected with CHIKV-ZsGreen, antiviral activity of screened compounds was assessed through automated numbering of fluorescent foci (Fig. 1). First, in the primary screening compounds in the PRB were tested at 50 and 10 μM. After the detection of fluorescent foci, the fixed cells were stained with crystal violet to estimate cytotoxicity. Those compounds that showed at 50 μM more than 60% reduction in the number of fluorescent foci forming units (fFFU) relative to control (VC, 1% DMSO) without presenting cytotoxicity (<10%) (n=20) and those compounds that, despite presenting cytotoxicity at 50 μM, showed >60% reduction in the number of fFFU and <50% cytotoxicity at 10 μM (n=16), were selected for secondary screening (Fig. 2a).

**Fig. 1.**
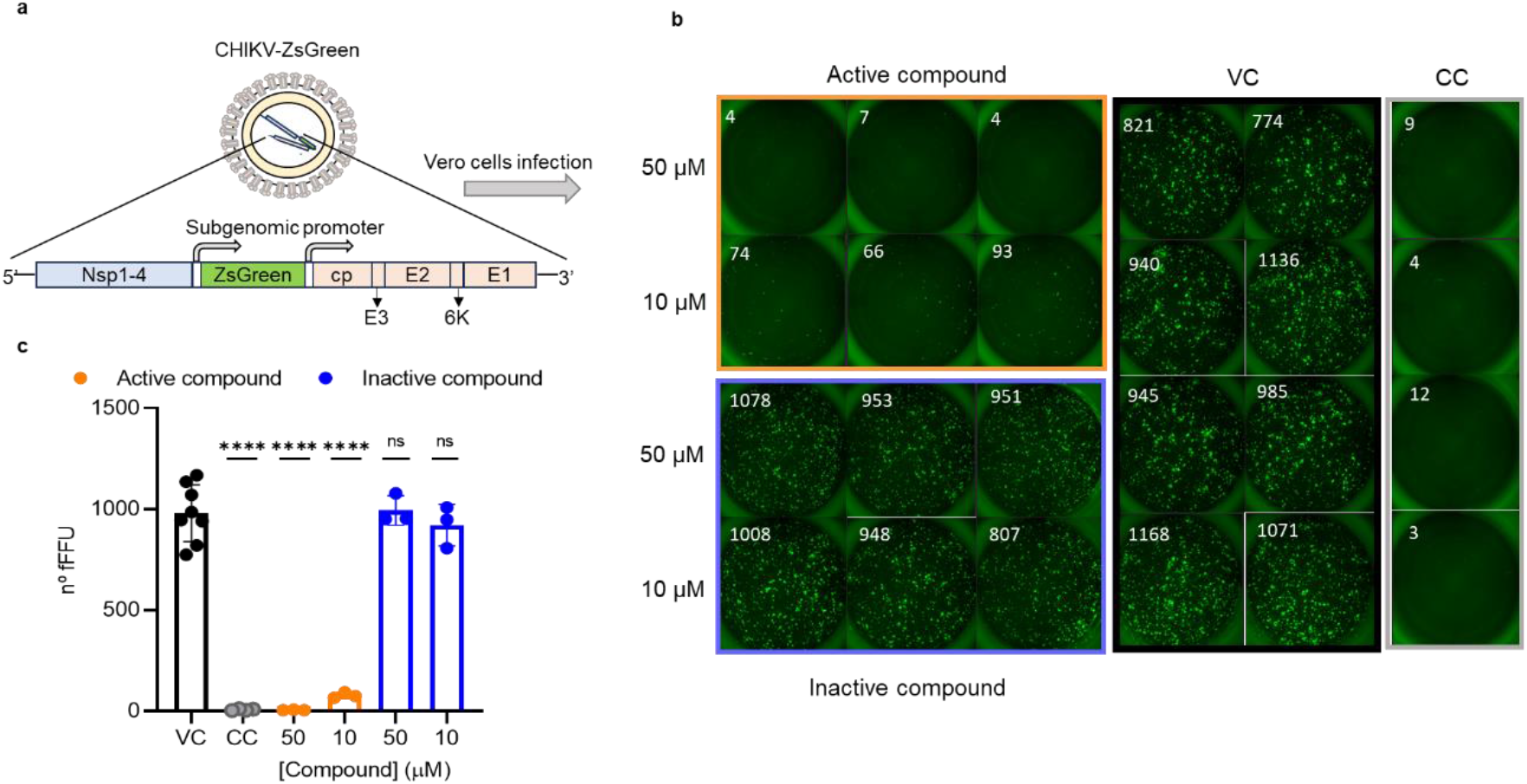
Reporter-based HTS antiviral assay using infectious CHIKV expressing ZsGreen (CHIKV-ZsGreen) (a) Schematic representation of CHIKV-ZsGreen reporter virus and its genome. When Vero cells are infected with CHIKV-ZsGreen, ZsGreen is expressed in the cell cytoplasm under the control of the virus subgenomic promoter. (b) Representative image of automated numbering (White numbers) of fluorescent infectious foci (fFFU) using the EliSpot equipment and the InmunoSpot 7.0.30.2 software (ImmunoSpot C.T.L, Germany) in the reporter-based HTS antiviral assay. For primary screening, Vero cells were infected with at least 700 FFUs of CHIKV-ZsGreen and simultaneously treated with 50 and 10 μM of the tested compounds in triplicate. Untreated infected cells (VC) and uninfected cells (CC) were included as controls. (c) Number of fFFU counted in (b). Unpaired t test vs VC: **** p< 0.0001, ns: not significant

**Fig. 2.**
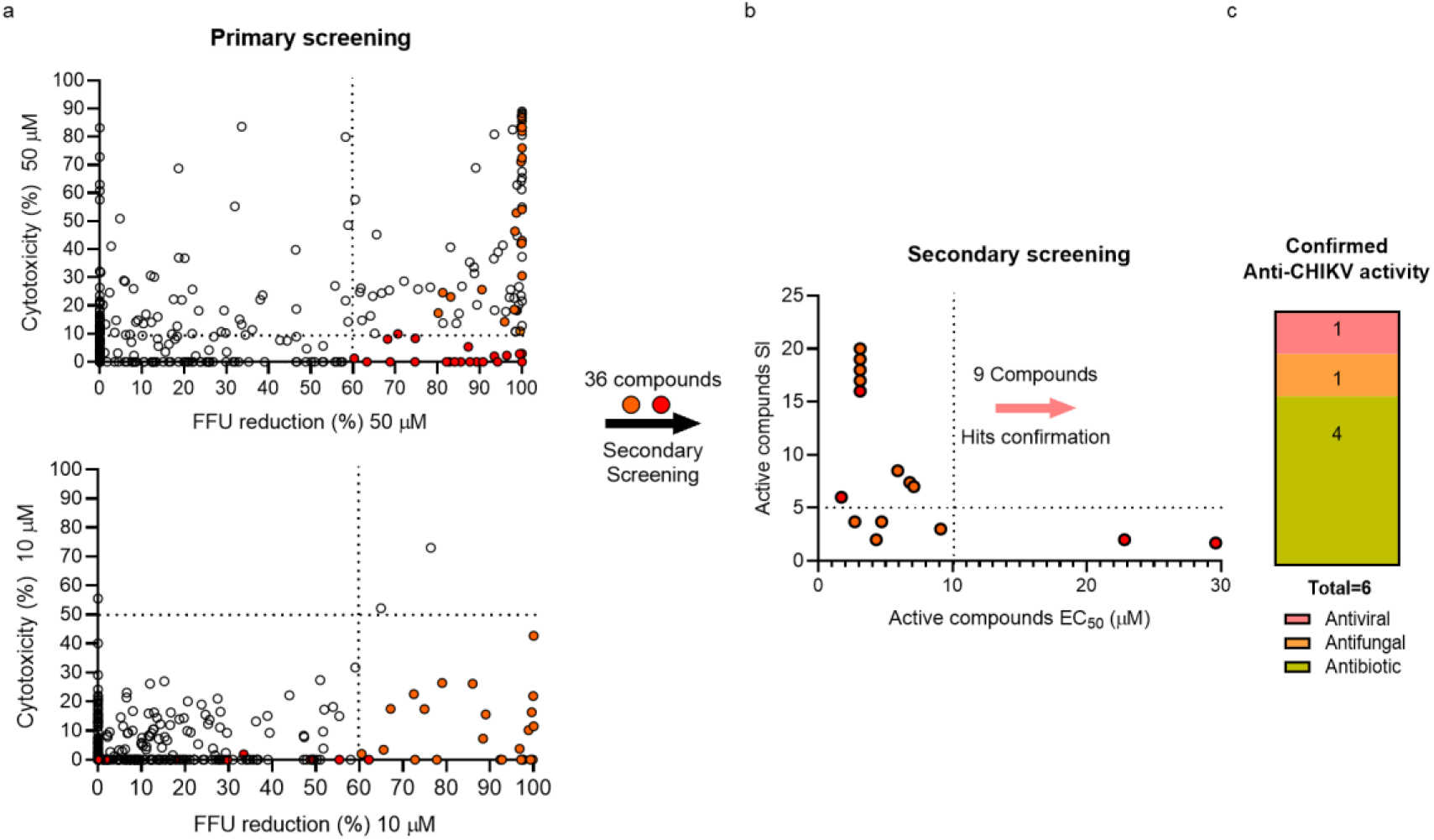
Identification of anti-CHIKV hits from Pandemic response box using infectious CHIKV-ZsGreen. (a) In the primary screening Vero cells were infected with CHIKV-ZsGreen and treated at compound concentrations of 50 and 10 μM. Cytotoxicity was estimated with crystal violet. Red dots indicate compounds that showed at 50 μM more than 60% reduction in the number of fluorescent foci forming units (fFFU) relative to mock treated control (VC) without presenting cytotoxicity (<10%) (n=20) and orange dots, those compounds that despite presenting cytotoxicity at 50 μM, showed >60% reduction in the number of fFFU and <50% cytotoxicity at 10 μM (n=16). (b) Secondary screening: compounds selected in the primary screening were tested in five serial dilutions and resulting active compounds are shown (n=15). Effective concentration 50 (EC_50_) values, which are defined as the concentration of compound that reduces fFFU number by 50% with respect to VC, were estimated from dose-response curves. Cytotoxicity was evaluated by MTS/PMS method. Selectivity index (SI) was calculated as: CC_50_/ EC_50_. Compounds with EC_50_ < 10 μM and SI >5 (n=9) were selected for hit confirmation assays. (c) Hit confirmation. EC_50_ values were estimated from dose-response curves and cytotoxicity was evaluated by MTS/PMS method in three independent assays.

Then, in the secondary screening, dose-response curves were performed starting at 50 or 10 μM to estimate the EC_50_ values for each compound. In parallel, CC_50_ was determined by MTS/PMS. Nine compounds (MMV1578574; MMV1578897; MMV1581545; MMV689401; MMV1634402; MMV102270; MMV000008; MMV690621; MMV1593535) showed fFFU reduction in a concentration-dependent manner with EC_50_ < 10 μM and SI >5 and were selected for hit confirmation assays (Fig. 2b). In turn, the reminder 27 compounds were cytotoxic at concentrations where antiviral activity was evidenced or presented poor SI (Online Resource 1) and were not further investigated.

### Hits confirmation: 4 new anti-CHIKV candidate drugs

Firstly, to confirm the antiviral activity and determine the selectivity of compounds emerging from screening, dose response curves against CHIKV-ZsGreen were performed and the cytotoxicity in Vero cells was assessed by MTS/PMS method. Results indicated that six out of the nine compounds evaluated showed potent antiviral activity, whereas three (MMV1593535, MMV1578897, and MMV1581545) were not active at non-cytotoxic concentrations or showed SI <5. From the six confirmed hits, one displays known activity as an antifungal agent (MMV689401), one as an antiviral agent (MMV690621) and four as antibiotics (MMV000008, MMV1578574; MMV1634402; MMV102270) (Fig. 2c). According to previous results, the antimalarial MMV000008 (Chloroquine) [22] and the broad spectrum indirect antiviral agent MMV690621 (GSK-983) [19, 23, 24] presented potent anti CHIKV activity and high selectivity, with EC_50_ values in the low micromolar and nanomolar range, respectively (Fig. 3). On the other hand, MMV689401, MMV1578574, MMV1634402, MMV102270 showed EC_50_ values in the low micromolar range (from 0.32 to 6.38 μM) and high SI (Fig. 3, Online Resource 1).

**Fig. 3.**
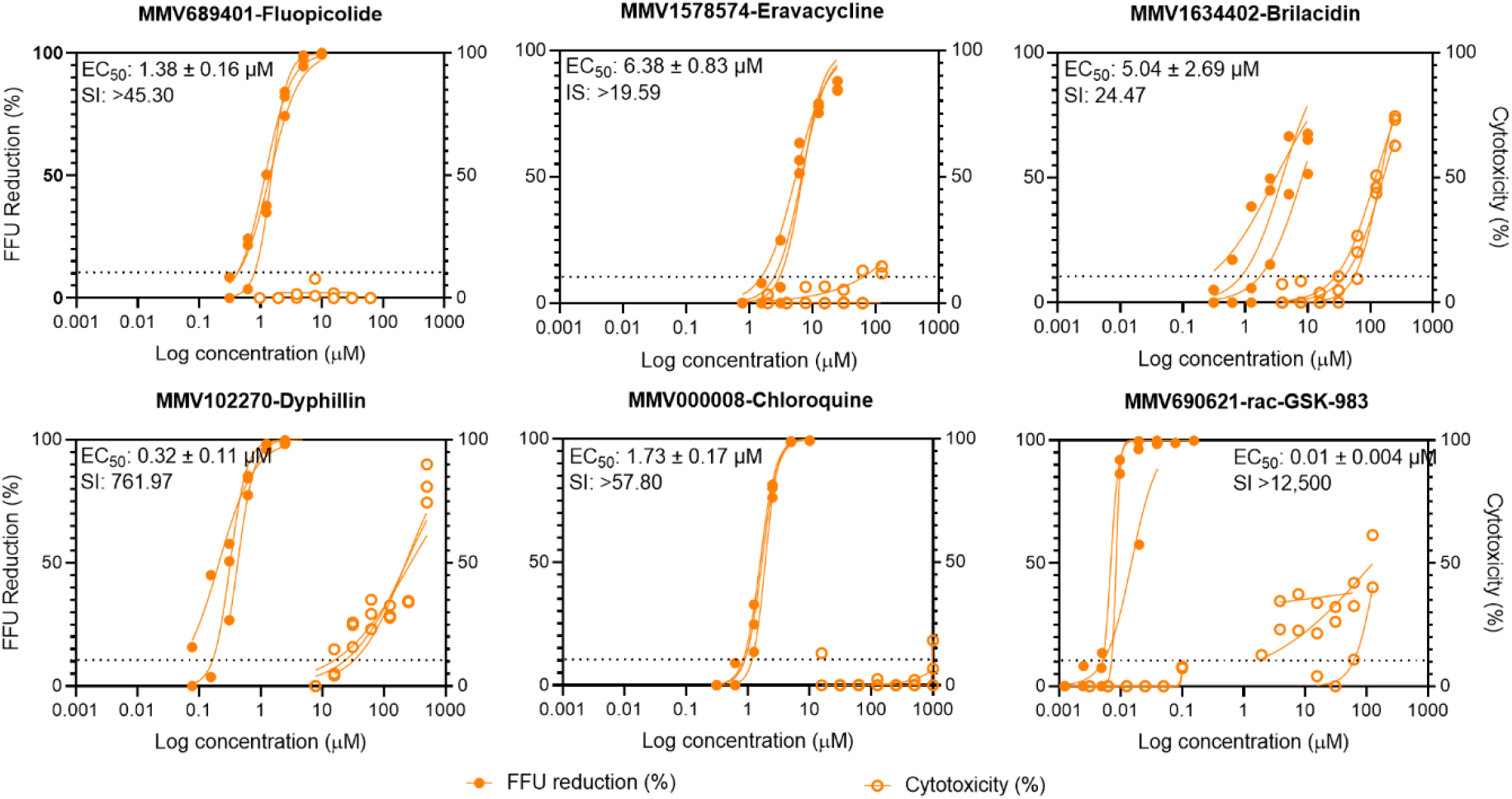
Anti-CHIKV activity and cytotoxicity of selected hits. To confirm the antiviral activity and determine the selectivity of compounds emerging from screening, dose response curves against CHIKV-ZsGreen (filled circles) were performed and the cytotoxicity in Vero cells (empty circles) was assessed by MTS/PMS method. Effective concentration 50 (EC_50_), which is defined as the concentration of compound that reduces fFFU number by 50% with respect to VC, and the cytotoxic concentration 50 (CC_50_), which is defined as the concentration of compound that reduces absorbance_490_ by 50% with respect to CC, were estimated from dose-response curves. Finally, the EC_50_ and CC_50_ values were used to calculate the Selectivity Index (SI = CC_50_/EC_50_)

Secondly, to verify the effect of active compounds on viral yield, culture supernatants from treated infected cells and VC were collected at 20 h p.i. and infectious virus particles were titrated by FFU. Results showed that the treatment with the identified hit compounds reduced the production of infectious progeny virus in a concentration-dependent manner (Fig. 4), adding new evidence on their anti-CHIKV activity. Compounds with EC_50_ in the low micromolar (MMV689401 and MMV000008) and nanomolar range (MMV102270 and MMV690621) achieved 100% fFFU reduction and showed viral yield titers approximately 100 times lower (98.6-99.8% reduction) than VC at the maximum non-cytotoxic concentration evaluated (MNCC). On the other hand, less potent compounds MMV1578574 and MMV1634402, achieved only 85.5 and 64.2% fFFU reduction at MNCC and viral yield titers were nearly 10 times lower (94.9 and 94.1% reduction) than VC.

**Fig. 4.**
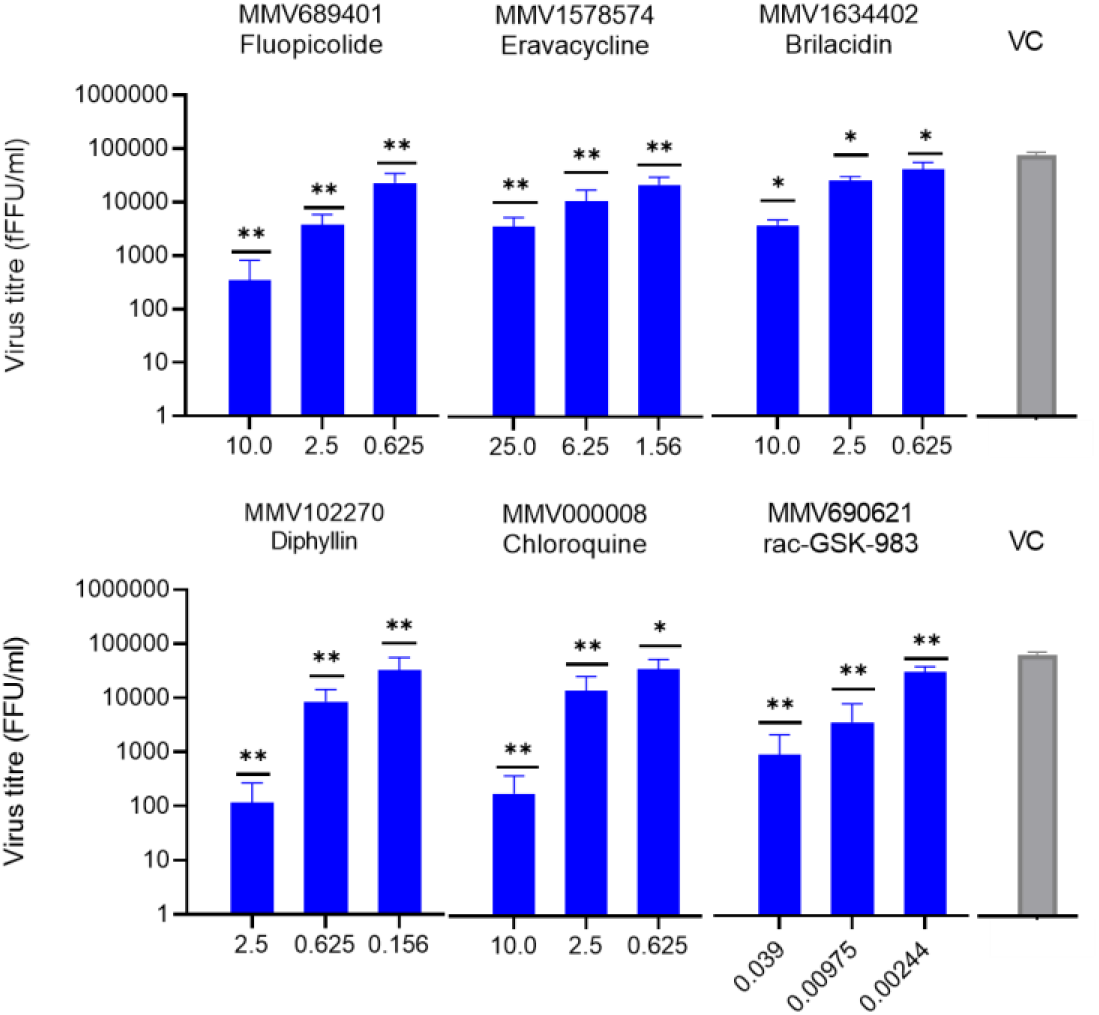
Viral yield assays. Culture supernatants collected from hits confirmation assays (Fig. 3) were titrated in Vero cells by FFU method. Mann-Whitney test vs VC *p<0.05; **p<0.005

## Discussion

The present work identified and validated 4 new molecules within the Pandemic Response Box as potential drug candidates against CHIKV: MMV1634402, MMV102270, MMV689401, and MMV1578574. In a previously reported screening of PRB compounds for antiviral activity against CHIKV, Policastro et al. [19] confirmed the activity of one compound which was identified following a screening strategy that used a cell line that stably expresses a CHIKV subgenomic replicon encoding viral proteins nsp1-nsp4 and a luciferase reporter gene. This system allowed sensitive identification of molecules that interfere with translation and genome amplification directed by non-coding regions and non-structural proteins. In the present work, we used a fully infectious CHIKV-ZsGreen reporter virus as a tool for screening, which reproduces the complete viral cycle and generates a productive infection in cultured Vero cells. This system allowed us to expand the range of potential targets to molecules that interfere with processes mediated by the structural proteins of the virus, including virus entry to the host cell, uncoating, morphogenesis and release of progeny viral particles. Accordingly, our screening approach successfully identified previously reported CHIKV antiviral compounds in the PRB and further expanded potentially active hits to compounds that presumably target steps in the replication cycle that involve the function of structural proteins. Among the compounds that resulted active in the primary and secondary screening, MMV690621 was previously reported to have anti-CHIKV activity in the subgenomic replicon-based assays reported by Policastro et al. [19], and MMV000008 (Chloroquine) was previously shown to inhibit virus entry and release [22]. Overall, our approach using fFFU reduction of CHIKV-ZsGreen as a screening method effectively identified compounds active against CHIKV.

Regarding the new hits identified here, the antiviral activity of MMV1634402 (Brilacidin) and MMV102270 (Diphyllin) against different viruses has been reported previously. Brilacidin is a synthetic, nonpeptidic, small molecule mimetic of defensin, a type of host defense proteins/peptides (HDPs) or antimicrobial peptides (AMPs), with a potent antibacterial activity against both Gram‐positive and Gram‐ negative bacteria [25]. It was evaluated in Phase 2 clinical trials both for bacterial and viral infections (Clinical Trials NCT02324335, NCT04784897 NCT02052388, NCT04240223, NCT01211470). Early reports on antiviral activity of brilacidin demonstrated its activity against SARS-CoV-2 [26, 27] and other human coronaviruses [28]. Mechanistic studies showed that brilacidin would block SARS-CoV-2 viral entry [27] and a dual antiviral mechanism of action was suggested, including virucidal activity and binding to coronavirus attachment factor heparan sulfate proteoglycans on the host cell surface [28]. In accordance with these mechanisms of action, brilacidin was recently proposed as potential antiviral against enveloped viruses [29]. Anderson et al. [29], demonstrated brilacidin antiviral activity against encephalitic alphaviruses VEEV and EEEV, arthritogenic alphavirus SINV and bunyavirus Rift Valley fever virus (RVFV), whereas the inhibitory activity was only modest in the context of non-enveloped virus.

Diphyllin, is an arylnaphthalene lignan lactone that can be isolated from many traditional medicinal plants. It has multiple bioactivities reported such as antitumor, anti-inflammatory, and antimicrobial activities (reviewed in [30]) and was found to inhibit the replication of many fungal, bacterial, and protozoan pathogens (reviewed in [31]). Diphyllin was originally reported to decrease vacuolar (H+)ATPase (or V-ATPase) activity and to neutralize the pH of lysosomes at nanomolar concentrations [31, 32]. Endosome acidification is essential for the cell entry of several viruses including CHIKV and, accordingly, diphyllin has also a broad-spectrum antiviral activity. It was found to be active against SARS-CoV-2 [31], tick-borne encephalitis virus (TBEV), West Nile virus (WNV), Zika virus (ZIKV), RVFV, rabies virus (RABV) [31], A and B influenza viruses, dengue virus (DENV-2) [32] and SINV [33]. In addition, diphyllin and its derivatives have also been shown to have other cellular targets such as topoisomerase IIα and induce or modulate intracellular processes (apoptosis, autophagy, late interferon-γ production, nitric oxide levels, TNF-α, and IL-12 production) which (alone or in combination) would explain why viruses that do not depend on endosome acidification, such as herpes simplex virus type 1 (HSV-1) [31], or Human Immunodeficiency virus (HIV) [34], are also inhibited by diphyllin. Similarly, we showed brilacidin and diphyllin were active against CHIKV in the low micromolar range, adding new evidence on its broad-spectrum potential.

On the other hand, MMV1578574 (Eravacycline) and MV689401 (Fluopicolide) have little or no precedent as antiviral agents. Eravacycline, is a synthetic fourth generation tetracycline that works via inhibition of the 30S ribosomal subunit and was initially approved in August 2018 to treat complicated intraabdominal infections [35]. It was reported to be active against SARS-CoV-2 targeting the viral main cysteine-protease 3CLpro [36] and, using computational approaches, it was suggested as a potential drug molecule against monkeypox DdRp [37].

Finally, Floupicolide is a fungicide first registered in China in 2005 which is effective against a wide range of Oomycete (Phycomycete) diseases in plants that impact on global agricultural production. It exhibits low toxicity to plants, fungi, and humans [38, 39] and its primary target was suggested to be vacuolar H+/-ATPase subunit. To the best of our knowledge, there are no reports of fluopicolide activity against human pathogens, neither viral nor bacterial, being this the first report on its potential antiviral activity.

### Conclusions

Overall, our screening approach allowed identifying additional active hits against CHIKV within the PRB. Our results add new evidence on the broad-spectrum antiviral activities of some molecules that may interfere with viral entry and uncoating such as Brilacidin and Diphyllin, incorporates a new potential target for the antiviral action of a molecule with a possible anti cysteine-protease activity (Eravacycline) and introduce new perspectives on the potential use of Fluopicolide as an antiviral agent.

Future work on the study of their effect on CHIKV replication cycle as well as the selection and characterization of resistant virus will allow to get some insight about their mode of action and molecular target.

## Supporting information

Online Resource 2

Online Resource 1

## Acknowledgments

The authors gratefully thank the Medicine for Malaria Ventures (MMV, www.mmv.org) and the Drugs for Neglected Diseases initiative (DNDi, www.dndi.org) for the Pandemic Response Box and supplying the compounds. CHIKV La Reunion infectious clone expressing ZsGreen was a kind gift of Kenneth A. Stapleford (Department of Microbiology, New York University School of Medicine).

## Data availability

All source data of this study are available within the paper and associated Online Resources. Any additional information related to the study is available from the corresponding author upon reasonable request.

## Authors contribution

EFC performed antiviral assays and data analysis. EFC and DEA contributed equally to conceptualization, experimental design, and manuscript writing.

