## Supplementary material for "New highly selective antivirals for Chikungunya Virus identified from the screening of a drug-like compound library": Online Resource 2

**Supplementary Table 2. Hits confirmation.** Cytotoxicity and antiviral activity of compounds selected after secondary screening.

| MMV ID | [1]$EC_{50}$ (µM) | [2]$CC_{50}$ (µM) | [3]SI |
|---|---|---|---|
| MMV1578574 | 6.38 (0.83) | >125.00 | >19.59 |
| MMV1578897 | 5.51 (1.15) | >62.50 | >11.34 |
| MMV1581545 | Inactive | 167.77 (14.39) | nd |
| MMV689401 | 1.38 (0.16) | >62.5 | >45.0 |
| MMV1634402 | 5.04 (2.69) | 143.50 (20.19) | 24.47 |
| MMV102270 | 0.32 (0.11) | 243.83 (25.84) | 761.97 |
| MMV000008 | 1.73 (0.17) | >100.00 | >57.80 |
| MMV690621 | 0.01 (0.004) | >125.00 | >12,500 |
| MMV1593535 | 8.84 (1.97) | 41.97 (3.29) | 4.74 |

Nine compounds selected after secondary screening were tested in six serial dilutions in triplicate against CHIKV-Zs-Green in Vero cells. The focus count data for each antiviral concentration and the VC were used to plot dose-response curves that were fitted by nonlinear regression using GraphPad Prism 8 software. In parallel, cytotoxicity was determined by MTS/PMS. Results shown the mean and standard deviation (ds) from three independent experiments.

[1]$EC_{50}$: Effective concentration 50 is defined as the concentration of compound that reduces fFFU number by 50% with respect to mock treated infected cells.

[2]$CC_{50}$: Cytotoxic concentration 50 is defined as the concentration of compound that reduces cell viability by 50% with respect to mock treated cells.

[3]SI: Selectivity index, where $SI=CC_{50}/EC_{50}$
