## Supplementary material for "New highly selective antivirals for Chikungunya Virus identified from the screening of a drug-like compound library": Online Resource 1

**Supplementary Table 1. Secondary screening.** Cytotoxicity and antiviral activity of compounds selected after primary screening.

| **MMV ID** | **[1]$EC_{50}$ (µM)** | **[2]$CC_{50}$ (µM)** | **[3]SI** |
|---|---|---|---|
| MMV1580853 | 22.8 | >50.0 | >2.0 |
| MMV1578578 | 20.4 | >50.0 | >2.0 |
| MMV394033 | 9.1 | 28.5 | 3.0 |
| MMV1782105 | 29.6 | >50 | >1.7 |
| MMV1634360 | 4.7 | 17.5 | 3.7 |
| **MMV1578574** | **5.9** | **>50.0** | **>8.5** |
| **MMV1578897** | **6.8** | **>50.0** | **>7.4** |
| **MMV1581545** | **7.1** | **>50.0** | **>7.0** |
| MMV233495 | 4,3 | >10.0 | >2.0 |
| **MMV689401** | **<3.1** | **>50.0** | **>16.0** |
| **MMV1634402** | **<3.1** | **>50.0** | **>16.0** |
| **MMV102270** | **<3.1** | **>50.0** | **>16.0** |
| **MMV000008** | **<3.1** | **>50.0** | **>16.0** |
| **MMV690621** | **<3.1** | **>50.0** | **>16.0** |
| **MMV1593535** | **1.7** | **>10.0** | **>6.0** |

For secondary screening, 36 compounds selected after primary screening were tested in five serial dilutions in triplicate against CHIKV-Zs-Green in Vero cells. The focus count data for each antiviral concentration and the VC were used to plot dose-response curves that were fitted by nonlinear regression using GraphPad Prism 8 software. In parallel, cytotoxicity was determined by MTS/PMS. Compounds with antiviral activity at non-cytotoxic concentrations are shown. The reminder 21 compounds were cytotoxic at concentrations where antiviral activity was evidenced and were not included. Compounds in bold were selected for further hits confirmation assays.

[1]$EC_{50}$: Effective concentration 50 is defined as the concentration of compound that reduces fFFU number by 50% with respect to mock treated infected cells.

[2]$CC_{50}$: Cytotoxic concentration 50 is defined as the concentration of compound that reduces cell viability by 50% with respect to mock treated cells.

[3]SI: Selectivity index, where $SI=CC_{50}/EC_{50}$
